# Farm productive realities and the dynamics of bovine viral diarrhea (BVD) transmission

**DOI:** 10.1101/230045

**Authors:** Bryan Iotti, Eugenio Valdano, Lara Savini, Luca Candeloro, Armando Giovannini, Sergio Rosati, Vittoria Colizza, Mario Giacobini

## Abstract

Bovine Viral Diarrhea (BVD) is a viral disease that affects cattle and that is endemic to many European countries. It has a markedly negative impact on the economy, through reduced milk production, abortions, and a shorter lifespan of the infected animals. Cows becoming infected during gestation may give birth to Persistently Infected (PI) calves, which remain highly infective throughout their life, due to the lack of immune response to the virus. As a result, they are the key driver of the persistence of the disease both at herd scale, and at the national level. In the latter case, the trade-driven movements of PIs, or gestating cows carrying PIs, are responsible for the spatial dispersion of BVD. Past modeling approaches to BVD transmission have either focused on within-herd or between-herd transmission. A comprehensive portrayal, however, targeting both the generation of PIs within a herd, and their displacement throughout the Country due to trade transactions, is still missing. We overcome this by designing a multiscale metapopulation model of the spatial transmission of BVD, accounting for both within-herd infection dynamics, and its spatial dispersion. We focus on Italy, a country where BVD is endemic and seroprevalence is very high. By integrating simple within-herd dynamics of PI generation, and the highly-resolved cattle movement dataset available, our model requires minimal arbitrary assumptions on its parameterization. Notwithstanding, it accurately captures the dynamics of the BVD epidemic, as demonstrated by the comparison with available prevalence data. We use our model to study the role of the different productive realities of the Italian market, and test possible intervention strategies aimed at prevalence reduction. We find that dairy farms are the main drivers of BVD persistence in Italy, and any control strategy targeting these farms would lead to significantly higher prevalence reduction, with respect to targeting other production compartments. Our multiscale metapopulation model is a simple yet effective tool for studying BVD dispersion and persistence at country level, and is a good instrument for testing targeted strategies aimed at the containment or elimination of this disease. Furthermore, it can readily be applied to any national market for which cattle movement data is available.

## Introduction

Bovine Viral Diarrhea (BVD) virus is a pathogen responsible for a livestock disease of major concern in Europe, causing high prevalence in affected countries and important economic impacts (Lindberg et al., 2006). Induced costs are mainly due to production losses, derived from the immunosuppressive and abortive actions of the etiological agent, and to the biosecurity and immunization measures often implemented for its control or eradication (Thomann et al., 2017; Lindberg et al., 2006; Gunn et al., 2004). In addition, the disease can also facilitate the introduction and spread of other pathogens (Servizio Informativo Veterinario, 2011).

BVD normally displays mild clinical symptoms, though it may predispose affected animals to more severe forms of enteric and respiratory ailments, and in some cases result in the highly lethal Mucosal Disease. Transmission mainly occurs perorally or nasopharingeally, but excretion in semen, as well as iatrogenic diffusion and environmental persistence have been reported (Lanyon et al., 2014; Niskanen and Lindberg, 2003; Traven et al., 1991; Kirkland et al., 1991; Meyling and Mikél Jensen, 1988). Pregnant cattle are the segment of the population that is most at risk, and of the greatest epidemiological relevance. If viral contact occurs during the very initial phase of gestation, abortion will likely occur. Contact during the first third of pregnancy has instead a high probability of resulting in an immunotolerant persistently infected (PI) calf. Infection during later stages of pregnancy can result in fetal malformations and a considerable number of abortions.

PIs are the main epidemiological agent responsible for disease diffusion and persistence within a farm. Their viral excretion is orders of magnitude larger than that of an immunocompetent transiently infected animal, and is lifelong (Ezanno et al., 2007). Pregnant dams that carry a PI are latent persistently infected animals (PILs) and hardly detectable by serological tests as they are immune (Lanyon et al., 2014). They are an epidemiologically important aspect of the infection dynamics as their offspring will introduce PIs in the farm, potentially leading to disease persistence. PI animals tend to be smaller and underdeveloped compared to their healthy peers. Their lifespan is considerably shorter and their productivity severely hampered, so that they are more likely to be sold, thus further increasing the spatial spreading potential.

No therapy exists for BVD other than symptomatic treatment. Vaccines are available, but their effectiveness is often limited by the extremely high infectious pressure of PI animals. For these reasons, the current approach to BVD prevention mainly revolves around the implementation of biosecurity measures aimed at preventing contacts between infected animals and pregnant cows (Courcoul and Ezanno, 2010; Lindberg et al., 2006; Ezanno et al., 2007; Lanyon et al., 2014; Ezanno et al., 2008). Large variations across countries are however observed, and efficiency of the implemented measures is hard to evaluate. Modeling thus provides a convenient framework to study BVD dynamics, identify key drivers for transmission and propose control and eradication strategies.

Much research has been devoted to the development of BVD models for within-farm spread (Viet et al., 2007; Ezanno et al., 2007; Gunn et al., 2004; Cherry et al., 1998; Innocent et al., 1997b,a; Sørensen et al., 1995; Damman et al., 2015; Smith et al., 2009; Ezanno et al., 2008; Viet et al., 2004). Models focused on the complexity of the infection dynamics (e.g. horizontal and vertical transmission, persistently infected and transiently infected animals) and its parametrization, the disease consequences on herd demography (e.g. impact on reproduction, probability of abortion, reduced lifespan), the role of herd structure and herd-management practices, the economic impact and the evaluation of interventions. Though the presence of a single PI in an infection-free herd was estimated to pose a considerable risk with potentially long-term consequences (Innocent et al., 1997a), reseeding events were found to be critical for the persistence of the disease (Viet et al., 2004; Damman et al., 2015; Smith et al., 2009). The external risk of BVD introduction was generally modeled through synthetic importation schemes, only effectively accounting for geography and trade. Courcoul and Ezanno first proposed a spatially explicit metapopulation approach to model the regional spread of BVD in a group of 100 dairy herds assimilated to patches (Courcoul and Ezanno, 2010; Ezanno et al., 2008). Trade movements were however synthetically modeled through a random network. As such, they did not capture the heterogeneities and temporal correlations observed in cattle mobility that were found to strongly impact epidemic dynamics (Bajardi et al., 2011; Valdano et al., 2015; Ensoy et al., 2013; Vernon and Keeling, 2009; Ezanno et al., 2006). A data-driven spatial approach was proposed by Tinsley *et al.* that integrated data on cattle movements between beef farms in Scotland to simulate BVD spread at a larger scale and identify movement-informed interventions (Tinsley et al., 2012). To account for the complexity intrinsic to the network of cattle trade movements, the infection dynamics was however simplified through the use of a susceptible-infected-susceptible process where farms are treated as single units with no further population substructure. Moreover, only farms of one productive type were considered.

The aim of this work is to propose a novel multiscale spatially explicit metapopulation approach to model the geographic dispersion of BVD at the national scale, accounting for within-farm dynamics and high-resolution cattle movement data. Heterogeneities of BVD incidence per farm and reintroduction events are the result of the explicitly modeled dynamics, self-consistently producing geotemporal estimates for spreading potential and probability of exposure without the need for synthetic assumptions. We apply our study to the cattle population of Italy and more specifically we focus on the role that different productive realities may have in the spreading dynamics. We consider the full animal-level Italian database of cattle trade movements, accounting for different premises types (e.g. farms, slaughterhouses, markets, assembly centers, etc.) and four farm productive classes (beef with and without on-premise reproduction, dairy or mixed). We simulate different transmission profiles and generate simulated BVD prevalences at endemic equilibrium. Our findings on the role of different productive realities are then used to inform efficient targeted interventions aimed at a considerable reduction of prevalence in the country.

## 2. Materials and Methods

### 2.1 Data

We use the complete dataset of all individual animal movements, births, deaths and thefts covering the period from January 1st to December 31st 2014 provided by the Istituto Zooprofilattico Sperimentale (IZS) dell’Abruzzo e del Molise, the Italian national reference centre for the bovine movement database. The dataset is similar to the ones analyzed in (Bajardi et al., 2011, 2012; Natale et al., 2011, 2009; Valdano et al., 2015).

For each movement, we have an anonymized animal ID along with the corresponding age, breed, sex, date of movement, origin and destination premises. We consider three distinct categories of animals: males, females below reproductive age, and females of reproductive age. For each premise, we have an anonymized node ID, structure and production type labels and initial values for each of the three animal categories. The percentage of females in each animal holding on January 1st, 2014 is also provided.

The bovine dataset covers a total of 5.9 million movements between 144,403 premises throughout the country, involving 3.9 million bovines during the year 2014 (Figure 1a). The movement dataset is represented in the form of a time-referenced directed weighted network, where nodes represent premises and links represent bovine movements (Bajardi et al., 2011, 2012; Natale et al., 2011, 2009; Valdano et al., 2015).

**Figure 1:**
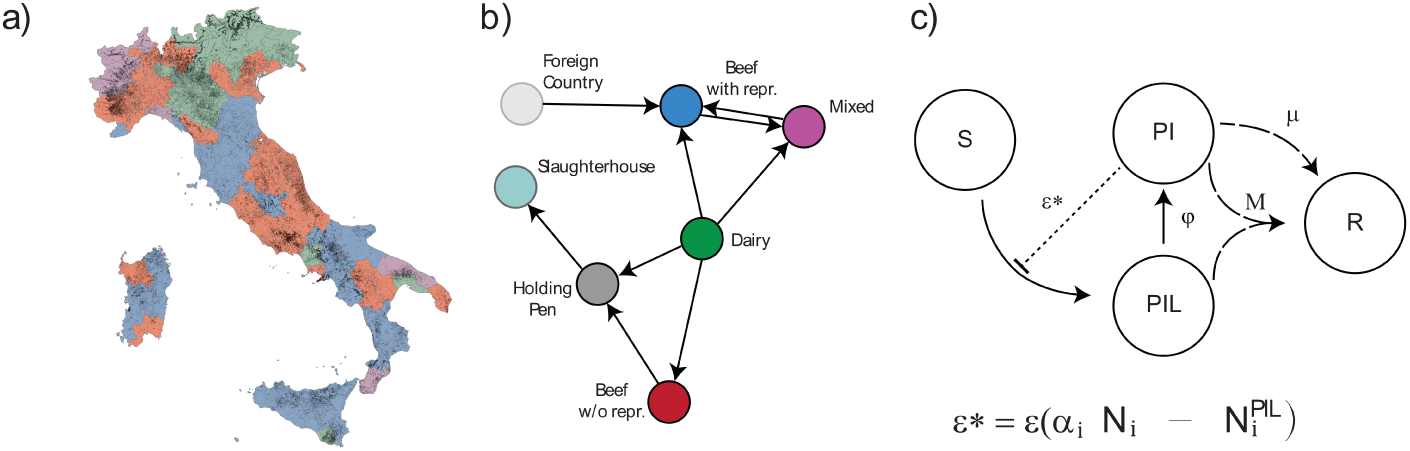
Metapopulation model for BVD transmission at the national scale. (a) Geographical visualization of the position of the 144,403 cattle premises in Italy of the 2014 movement database. Provinces are color coded according to the productive class mostly present in the area (red for beef fattening farms, blue for beef reproductive farms, green for dairy and violet for mixed attitude farms, as in panel (b)). Farms are represented as points on the map. (b) Schematic representation of the network of cattle movements where animal premises are labeled according to their type (farm, slaughterhouse, etc.) and productive class (beef with and without reproduction, dairy, mixed). (c) Scheme of the infection dynamics within each premise.

### 2.2. Model

We propose a multiscale spatially explicit metapopulation approach that simulates disease transmission between animals within each farm and disease dispersal in space through the movement of infected animals. Animal premises correspond to the patches of the model where BVD infection dynamics occurs, and trade movements correspond to the spatial coupling between patches, bringing the description of the disease spread from a local perspective to the national scale.

The infection dynamics is based on a compartmental model that represents the BVD spread in cattle herds, and includes the persistently infected animals (PI) and the persistently infected (latent) animals (PIL) (Figure 1c). A PI can infect a susceptible cow (S) and generate a PIL with probability *ε*\* given by the transmission rate *ε* times the number of susceptible female cows within the farm:

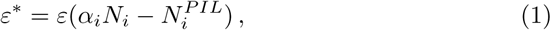

where *N_i_* is the total number of animals within premise *i*, 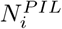 is the number of PIL animals within the same premise, and *α_i_* is the fraction of females present at that time in the premise. To account for the gestation stages and birth events, a PIL has then a probability *φ* of becoming a PI, expressed as 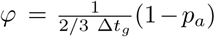, where Δ*t_g_* represents the average cattle gestation length and *p_a_* the probability of having an abortion. PIs are removed with rate *μ* accounting for both the infection-induced mortality and the culling of detected infected animals. In addition, rate *M* corresponds to the possibility of developing the Mucosal Disease, after which the farm will be cleared of all PI and PIL animals through death or abortion. The parameter *M* is also the only direct route for elimination of PILs, as they are unaffected by the processes modeled through *μ* (i.e. they are not clinically affected by the disease and are hard to detect).

The infection dynamics are applied to all animal holdings. Heterogeneity in within-farm disease dynamics is given by the farm productive class and associated herd management and demography. Here we consider four different productive types of interest (Figure 1b), similarly to (Dutta et al., 2014; Natale et al., 2009):

1. Dairy: all farms concerned with dairy production;
2. Beef with reproduction: farms hosting beef cattle with on-premise reproduction;
3. Beef without reproduction: farms fattening and raising beef cattle but not having on-premise reproduction;
4. Mixed: all farms that cannot be ascribed to a single productive type.

Cattle movements between premises are responsible for the spatial dispersion of a BVD epidemic. These are sold and purchased from farm to farm or through markets and may be divided into three categories of male cattle, and female cattle below or above reproductive age. Within each demographic category, animals are moved according to trade data and the probability that a moving animal belongs to a given disease status is simply proportional to the fraction of animals in that disease class. Only females at or above reproductive age may move a PIL, while all classes may move a PI. Newborns and animals imported from foreign countries are assumed to be healthy and susceptible.

All transitions (related to the infection dynamics or the movements) are modeled as discrete and stochastic processes to account for stochastic fluctuations due to small population sizes inside premises. The time step of the simulation is 1 week. The model is implemented in Scala using the Breeze libraries for Mathematics. Each premise uses a thread-local implementation of the Mersenne-Twister random number generator from the Apache Commons Math 3 library.

### 2.3. Model parametrization

The only free parameters of our model are the transmission rate *ε* and the removal rate *μ*. All other parameters are fixed and taken either from the literature or from expert opinion. Reproductive age is set at 15 months with a gestation length Δ*t_g_* = 36 weeks. Both values are chosen as intermediate values between the reproductive performances of dairy and beef breeds based on expert opinion. We consider a probability of abortions *p_a_* = 0.7 (Damman et al., 2015; Ezanno et al., 2007), thus yielding *φ* = 0.0125. The percentage *α_i_* of females in animal holding *i* is set from data at the beginning of the simulations. Development of the Mucosal Disease is considered to occur after an average period of 2.5 years (i.e. 130 weeks), similarly to (Kelling, 2004; Innocent et al., 1997a), thus yielding the rate 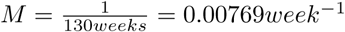.

Simulations start on January 1, 2014 and all cattle population is initialized in the susceptible class, except for a uniform seeding of 1% of all farms chosen at random. These farms are seeded with one PI each. This initial prevalence value was chosen as the lower estimate reported by Houe et al. (Houe, 1999) for the United Kingdom. The aim of our work is to study the equilibrium dynamics of the BVD epidemic, so we consider simulations running for several years on the 2014 weekly aggregated demographic and movement data with periodic boundary conditions. We use 1 year of data, as trade was rather stable in 2014, to neglect intrinsic changes over time due to the trade system and industrial expansions or reductions that can alter the equilibrium dynamics.

Removal rate is set to *μ* = 0.01*week*^−1^, and three different values of the transmission rate are explored − *ε* = *μ*/3, *ε* = *μ*, *ε* = 2*μ*. Results were computed over 50 different random seeding conditions for each transmission scenario explored.

### 2.4. Experimental scenarios of interventions

To support the findings of our study on the equilibrium dynamics of BVD circulation in Italy, we propose a set of experimental scenarios modeling interventions. They are aimed at isolating farms from the system, thus synthetically simulating the effect of 100% effective surveillance and control measures at the local scale. Assuming limited resources, we test removal strategies on a set of farms chosen with different approaches:

- removal of randomly chosen farms;
- removal of beef fattening farms without reproduction;
- removal of dairy farms;
- removal of mixed attitude farms.

The last three scenarios propose the targeted removal of farms based on their productive type thus accounting for their role in the epidemic dynamics, as emerged from our analysis. For each percentage value of farms removed, the targeted experimental scenarios are compared to the random removal, i.e. a choice not informed by the resulting epidemic relevance of each productive reality.

We explored removals of 1% to 24% of farms, in increments of 6%, which represents the maximum number of dairy nodes in the system. This is applied to the first three scenarios. Mixed farms are removed up to 12%, which is the maximum number of premises of that type, in increments of 6%. Each experimental scenario was repeated 10 times to stabilize model outputs.

### 2.5. Analyses

First, we analyze the Italian cattle trade dataset for 2014 to characterize the different productive realities in terms of farm size, trading patterns, and sex of animals.

Second, we observe the evolution in time of BVD prevalence in the cattle trade system, from the initial seeding condition to the equilibrium dynamics. Third, we explore the dependence of the simulated BVD prevalence on the transmission rate and on the productive realities. We consider different measures of prevalence, at the animal level and at the farm level, accounting for the productive type. More precisely, the local PI animal prevalence in farms of productive class *c* is computed as 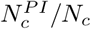,where 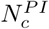 is the total number of PI animals in farms of productive class *c*, and *N_c_* is the number of these farms. We also compute the contribution of PI in farms of productive class *c* over all farms, to account for the sharply different sizes of the productive categories: 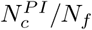, where *N_f_* is the total number of heads in all farms in the country. The national PI animal prevalence across all productive types is computed as 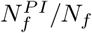, where 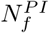 counts the total number of PIs in all farms. At the farm level, we consider a farm to be infected if there is at least one PI (or PIL). The national PI farm prevalence is therefore given by 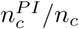 and by 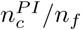, where 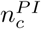 represents the number of PI infected farms of productive class *c*, *n_c_* is the number of farms of productive class *c*, and *n_f_* is the total number of farms in the country. Analogous measures were defined also for PILs.

Fourth, we assess the impact of the proposed experimental scenarios of interventions by comparing the resulting BVD national farm prevalence at equilibrium after the removal of a certain fraction of farms.

For each model output, average and 95% confidence interval (95% CI) are computed.

## 3. Results

Beef farms represent the most common productive reality in the Italian cattle trade system during the year 2014, with a total of 44,170 fattening farms (33.58% of all farms) and 36,350 reproduction farms (27.64%), amounting to more than 61% of the total number of farms (Figure 2a). They tend to be of small size, with a median of 3 heads (95% percentile 120) in fattening farms and 7 heads (95% percentile 69) in the reproduction ones (Figure 2b). Mixed production farms have a size similar to beef farms with reproduction (median 7 heads, 95% percentile 82), however they represent the smallest productive reality in our dataset (18,638 farms, 14.17%). Finally, dairy farms represent 24% of the system (32374 farms) and on average host the largest number of heads (median 23 heads, 95% percentile 207). The presence of females varies substantially across productive realities, with the largest values reported for milk production (average 55% of females according to data for January 1, 2014), reproduction purposes (41%) and mixed production (38%), and the smallest presence in beef farms without reproduction (14%).

**Figure 2:**
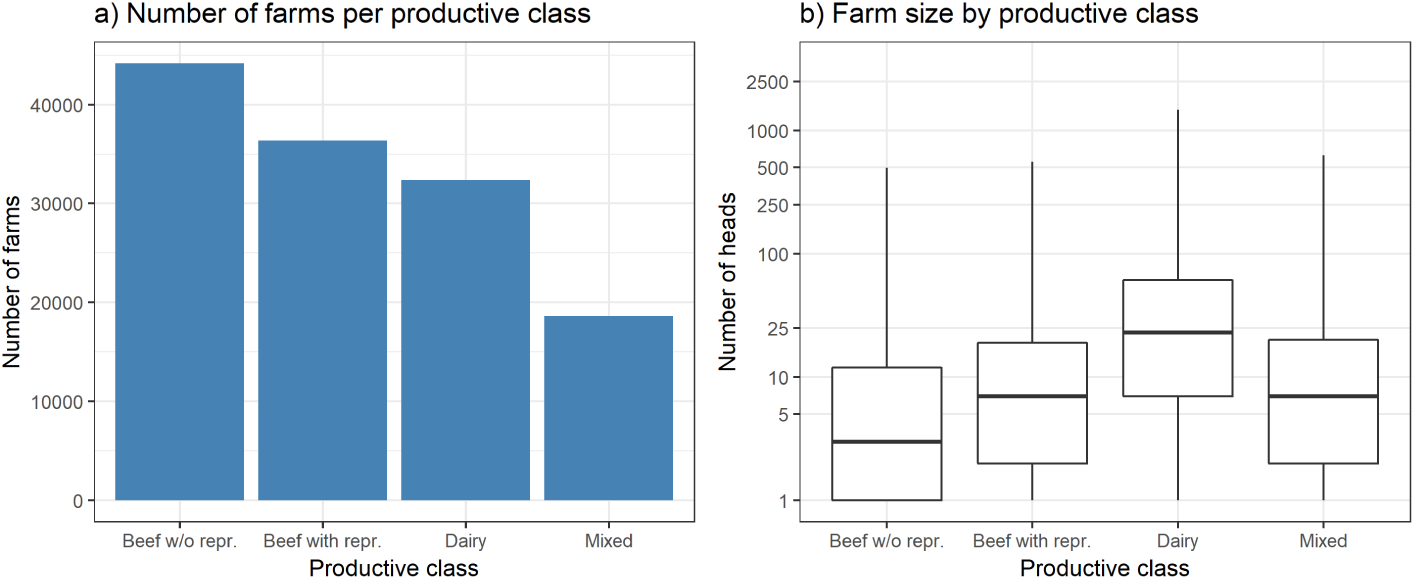
Description of productive realities in Italy in 2014. (a) Number of farms per productive class. (b) Distribution of the farm size per productive class. Wiskers are calculated as *median* ± 1.5 *IQR*

Trading patterns too are largely affected by production classes, with average trading volumes varying widely (Figure 3a). Dairy farms represent the most common origin of animal movements (44.7% of all movements), whereas beef farms without reproduction is the most common destination (54.3%). Movements follow a seasonal behavior, already outlined in (Bajardi et al., 2011; Valdano et al., 2015), with a considerable drop in activity around summer when animals are moved to pastures (these movements are not traced in the national database). Distinct trading patterns are observed when tracing the movements of female vs. male animals. A large number of male calves born every year in dairy farms are continuously sold to beef fattening farms before being moved to slaughterhouses at a later time (Figure 3b). The largest proportion of movements to slaughterhouses (65%) is indeed performed by beef fattening farms. A similar pattern is followed by beef farms with in-house reproduction. Female cows are strongly traded across dairy farms throughout the year.

**Figure 3:**
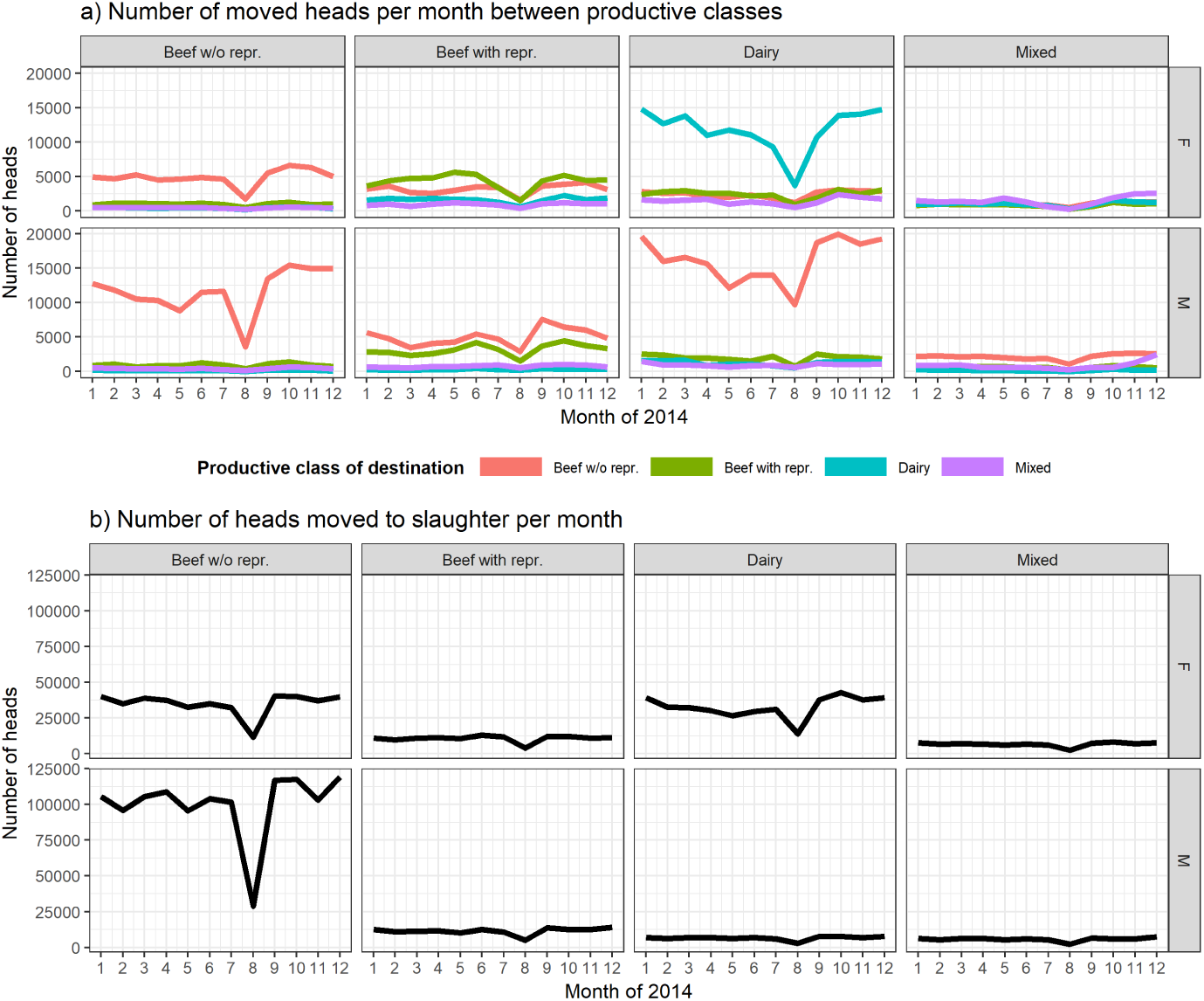
Movements of the 2014 cattle trade network in Italy. (a) Number of moved heads per month between productive classes. The title of each column correspond to the productive class of origin. Different colors refer to the productive class of destination. (b) Number of heads sent to slaughter each month. The title of each column correspond to the productive class of origin. In both panels, the top row reports the movements of female heads, the bottom row the ones of male heads.

This heterogeneous picture dependent on productive realities and affecting farm size, trading pattern and percentage of females becomes particularly relevant when modeling the spread of BVD and accounting for horizontal and vertical transmission as well as herd composition. Under the transmission conditions considered, our simulations reach the endemic equilibrium after about 20 years if *ε* > *μ* and after about 30 years if *ε* = *μ* (Figure 4). When *ε* < **μ**, the system maintains very low levels of prevalence, but the disease still persists in the cattle population. Interestingly, the described heterogeneities cause animal prevalence curves at the farm level to rapidly differentiate over time until they reach different values of BVD circulation at the endemic equilibrium. Mixed farms are the ones reporting the largest percentage of PIs, independently of the transmission condition, followed by dairy farms (Fig. 4a). PI animal prevalence within the productive class is equal to 3.54% (95% CI 3.51-3.58%) for mixed farms and to 3.027% (95% CI 3.018-3.036%) for dairy farms when *ε* > **μ** (1.73% (95% CI 1.72-1.76%) and 1.524% (95% CI 1.517-1.531%) for *ε* = **μ**, respectively). Dairy farms are however the ones contributing the largest amount of PIs to the system (Fig. 4b), leading to a predicted overall animal prevalence of 1.271% (95% CI 1.267-1.272%) for the higher transmission condition, and of 0.640% (95% CI 0.637-0.643%) for the intermediate value. This is also due to the large size of dairy farms compared to the other reproductive types.

**Figure 4:**
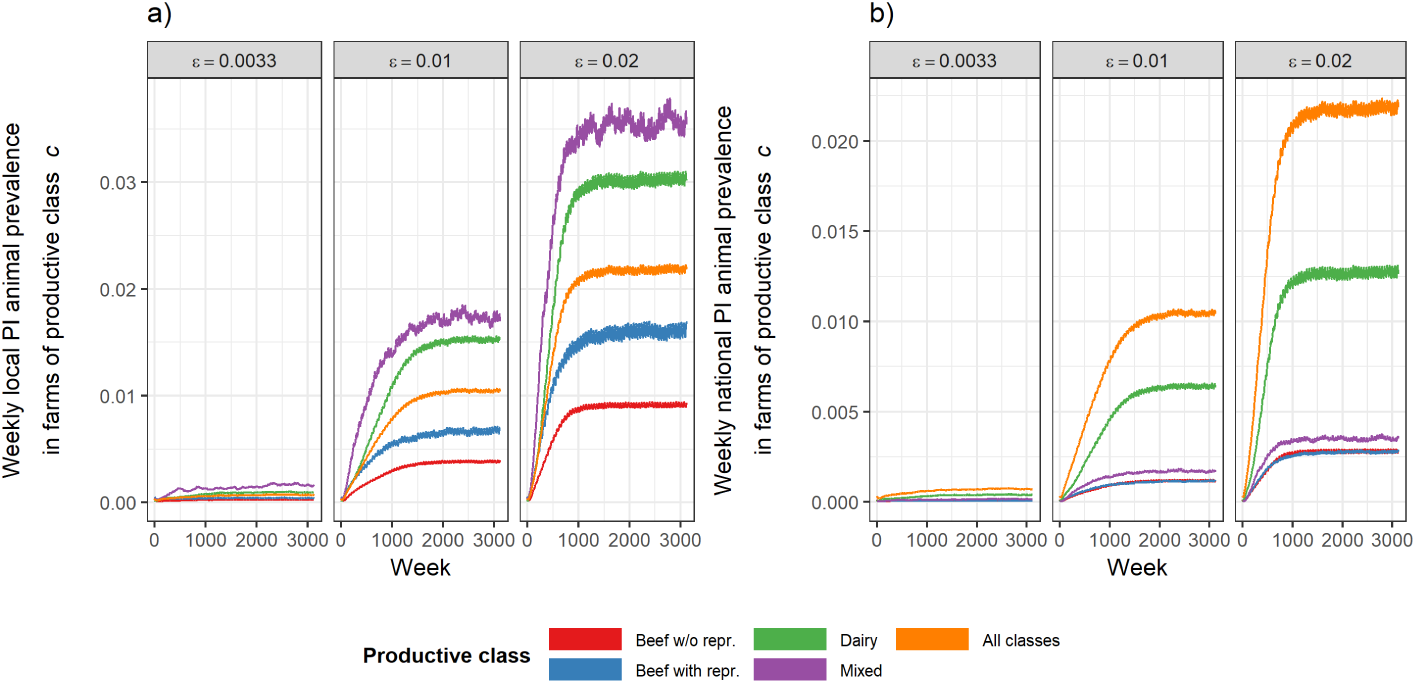
Weekly PI animal prevalence over time. (a) Average weekly local PI animal prevalence in farms of productive class *c*,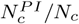. (b) Average weekly national PI animal prevalence in farms of productive class *c*, 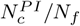. Each plot corresponds to a different value of the transmission rate ε. Colors refer to different productive classes or to the ensemble of all farms. Confidence intervals are small and not shown.

Figure 5 clearly shows that the ranking of productive realities that are mostly affected by the disease is maintained across transmission conditions. Most importantly, the overall contribution of PIs to the system by mixed and beef farms represent only 71.6% (95% CI 71.3-71.8%) of the contribution by dairy farms, indicating that the latter productive type dominates the circulation of BVD. Beef farms without on-site reproduction are the least affected. A picture similar to PI circulation is also observed for the spread of PILs, with the notable difference that within-productive class animal prevalence is higher in dairy farms with respect to mixed farms, though the two classes reach similar values at equilibrium (5.41% (95% CI 5.40-5.42%) vs. 4.99% (95% CI 4.97-5.01%), as shown in Figure 6). Since the generation of new PILs happens mostly in categories with high reproductive capability, beef farms with in-house reproduction show a higher prevalence in the case of PILs (up to 0.486%, 95% CI 0.484-0.488%) than fattening farms (up to 0.122%, 95% CI 0.122-0.123%). Finally we note that PILs reach a higher simulated prevalence than PIs, at 3.372% (95% CI 3.366-3.378%) vs 2.181 (95% CI 2.176-2.186%) in the high transmission condition.

**Figure 5:**
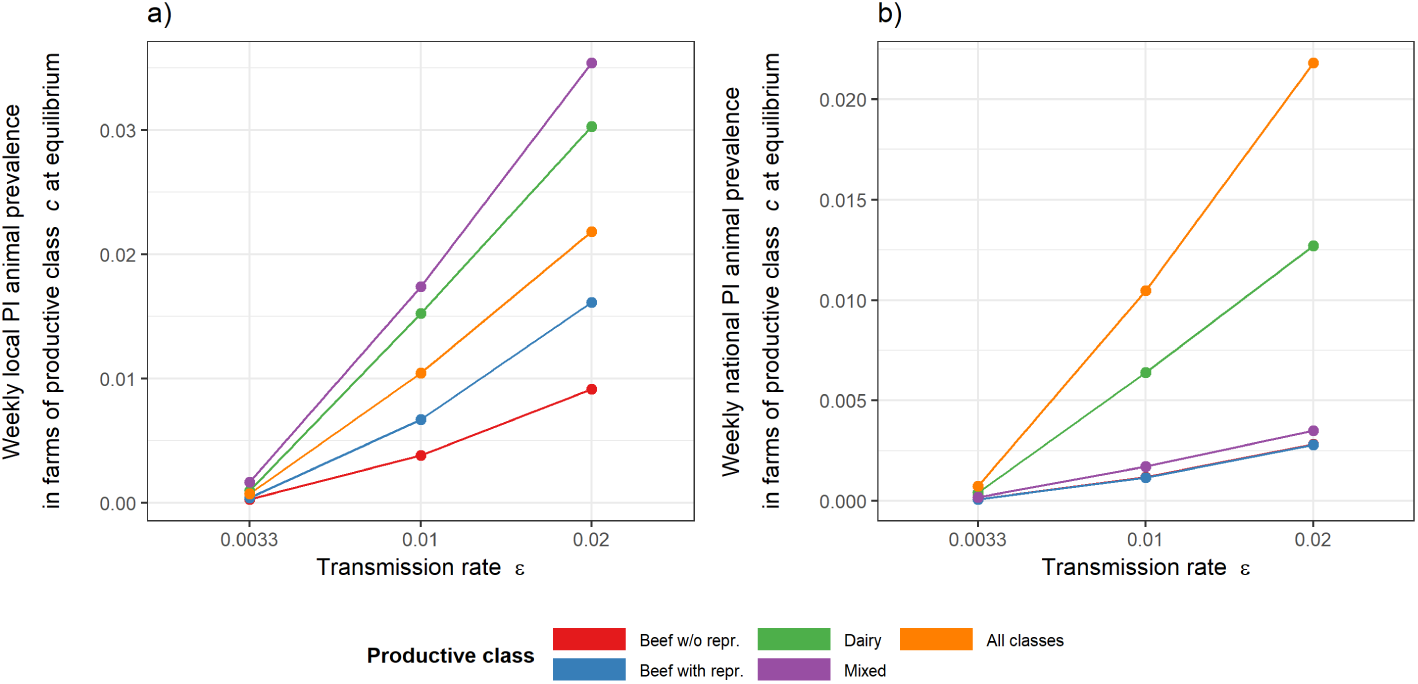
Weekly PI animal prevalence at endemic equilibrium. (a) Average weekly local PI animal prevalence in farms of productive class *c*, 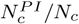, at endemic equilibrium as a function of the transmission rate *ε*. (b) Average weekly national PI animal prevalence in farms of productive class *c*, 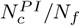, at endemic equilibrium as a function of the transmission rate *ε*. Colors refer to different productive classes or to the ensemble of all farms. Confidence intervals are small and not shown.

**Figure 6:**
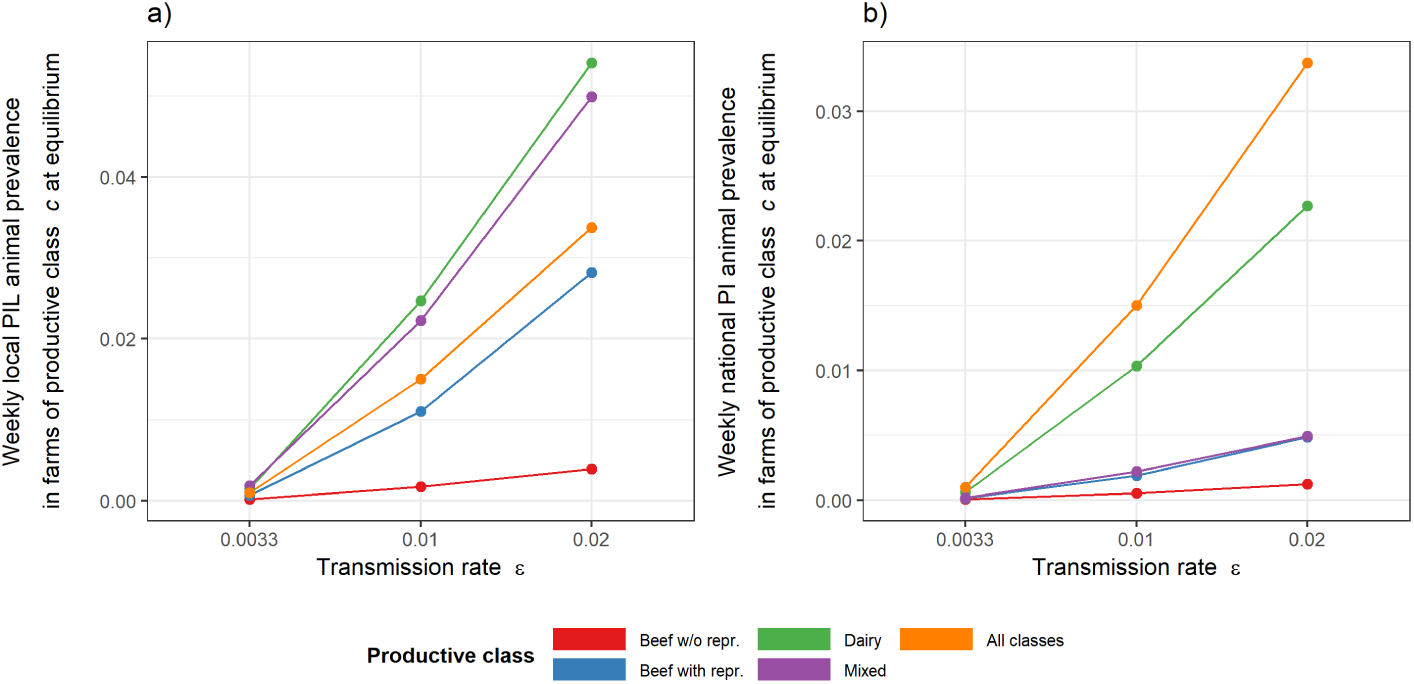
Weekly PIL animal prevalence at endemic equilibrium. (a) Average weekly local PIL animal prevalence in farms of productive class *c*, 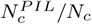, at endemic equilibrium as a function of the transmission rate *ε*. (b) Average weekly national PIL animal prevalence in farms of productive class *c*, 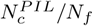, at endemic equilibrium as a function of the transmission rate *ε*. Colors refer to different productive classes or to the ensemble of all farms. Confidence intervals are small and not shown.

Dairy productive class results to be the most affected productive reality also when measuring the percentage of infected farms, with a predicted within-class prevalence of 13.96% (95% CI 13.94-13.98%) and across-class prevalence of 3.43% (95% CI 3.431-3.441%) (Figure 7). Beef fattening farms appear more relevant in contributing to the national farm prevalence than observed before for animal prevalence, likely because of the large number of smaller premises.

**Figure 7:**
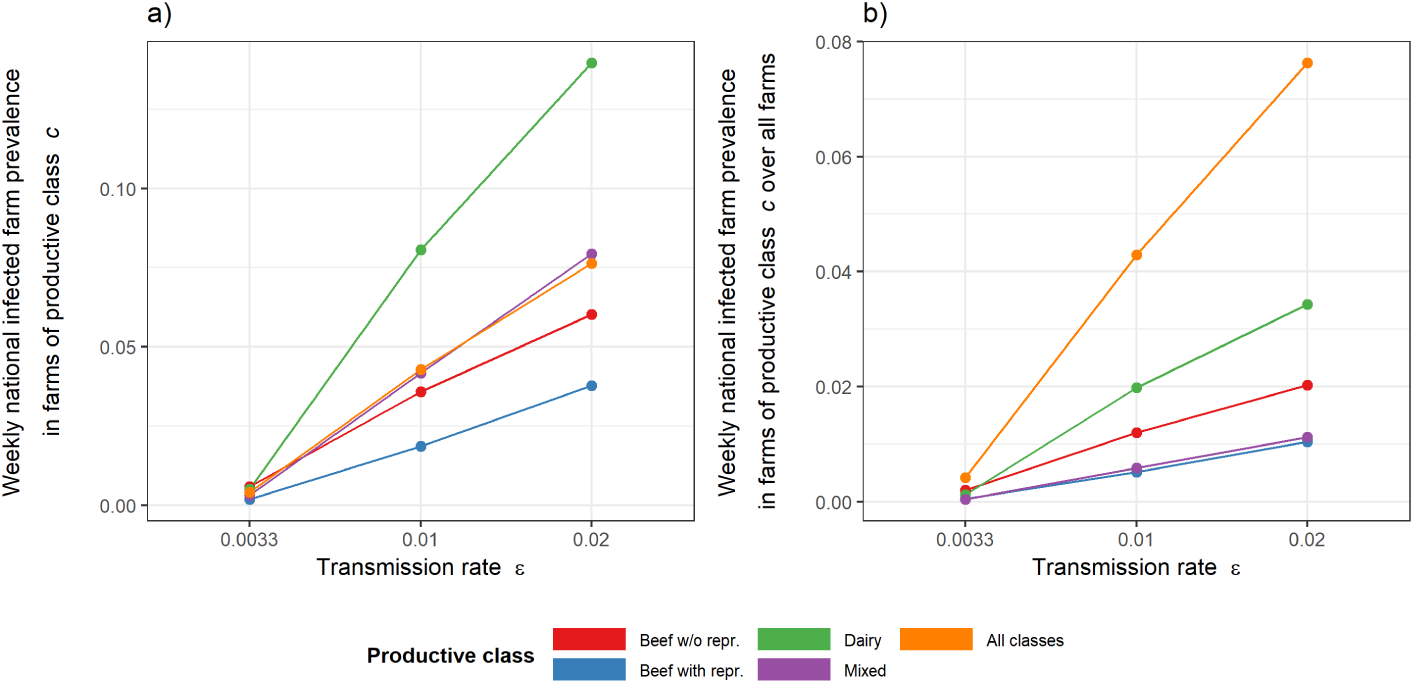
Weekly PI farm prevalence at endemic equilibrium. (a) Average weekly national PI farm prevalence in farms of productive class *c*, 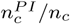, at endemic equilibrium as a function of the transmission rate *ε*. (b) Average weekly national PI farm prevalence in farms of productive class c over all farms, 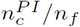, at endemic equilibrium as a function of the transmission rate *ε*. Colors refer to different productive classes or to the ensemble of all farms. Confidence intervals are small and not shown.

Given the central role of productive realities in the simulated endemic persistence of BVD in the cattle population in Italy, we test different experimental scenarios for intervention, comparing control measures randomly applied to farms with targeted ones based on their productive class. A dairy-focused intervention is predicted to be considerably more efficient in protecting the cattle population than a random farm intervention, for any degree of application of the intervention (Figure 8). Once 24% of the farms are removed, a dairy-targeted intervention leads to a drop of 78.8% (95% CI 78.7-78.9%) of the farm prevalence compared to the 46.1% (95% CI 45.9-46.3%) drop observed in the case farms were chosen randomly. Control measures focused on other productive classes are found instead to be less efficient than random interventions, as shown for example by the predicted 21.1% (95% CI 20.9-21.3%) drop when all beef fattening farms are targeted and the 22.7% (95% CI 22.4-23.0%) decrease when all mixed farms are selected.

**Figure 8:**
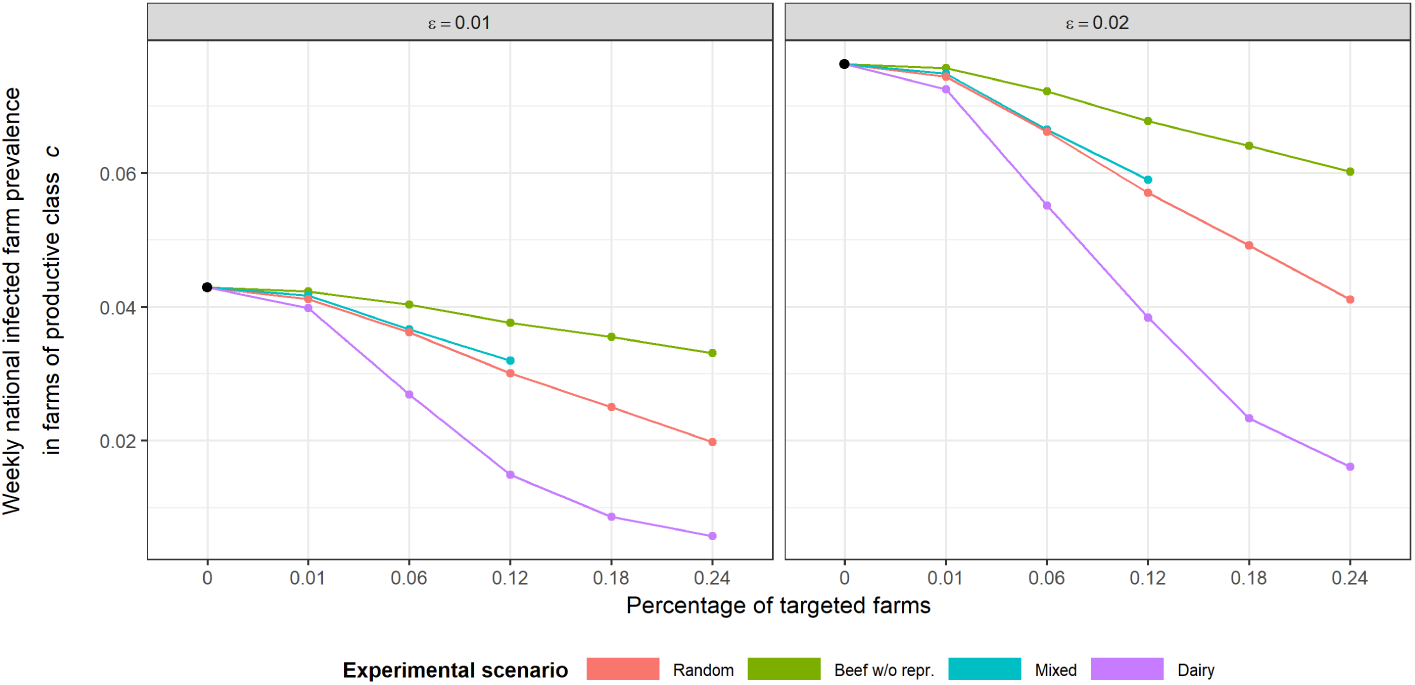
Random vs. targeted interventions. (a) Average weekly national infected farm prevalence in farms of productive class *c*, 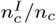, at endemic equilibrium as a function of the percentage of farms affected by the intervention. A farm is infected when it contains at least 1 PI or 1 PIL. Transmission rate is set to *ε* = 0.01. (b) as (a) for *ε* = 0.02. Different colors refer to different experimental scenarios for interventions. Confidence intervals are small and not shown.

## 4. Discussion

We use a spatially explicit metapopulation approach to model BVD endemic dynamics in the cattle population in Italy, based on demographic and mobility data for the year 2014. We find that farm productive realities strongly impact the circulation of the disease in the country, due to the specific sizes, herd structures and trading patterns associated to different productions. Dairy farms generate the largest number of persistently infected (PI) and latent persistently infected (PIL) animals. Basing their production mainly on self-sustenance through raising their own offspring internally, these farms represent the ideal habitat for a self-perpetuating and persistent BVD virus infection. Calves are indeed more often in close proximity with pregnant dams, present on the farm throughout the year, and this effect may in reality be further amplified by the fact that PIs are probably sold with preference because of reduced growth or insufficient productivity. As they also dominate national disease prevalence in the number of infected premises, we suggest dairy farms to be the ideal target for limited-resources control and interventions. Our results predict that dairy-focused measures would be almost twice as effective in prevalence reduction as measures applied randomly to the same number of farms independently of their productive reality. While representing a synthetic scenario for intervention, our proposed model could translate in the implementation of accurate and efficient local surveillance and biosecurity measures within a specific productive class, or immunization practices where available. For example, in areas where the disease is present and vaccination is not implemented, control strategies based on the analysis of bulk milk at dairy farms would greatly reduce the overall costs of testing while ensuring a good coverage and efficiency at the system level, as it has been done for other diseases (Muratore et al., 2017). In Switzerland, near-eradication has been achieved by serological testing of every head, followed by bulk milk surveillance of disease-free farms (Thomann et al., 2017). This experience further supports our model predictions and suggests a possible implementation in Italy as well. Given the importance of trade movements, quarantine measures following movements and the regulation of sales according to the disease status of a farm or a region would also be extremely important in BVD mitigation, as previously also suggested by (Courcoul and Ezanno, 2010) and as done for other diseases.

Other productive classes have a more limited role in the spread and maintenance of BVD. Fattening farms, for example, tend to be receivers and gatherers of PI animals. However these are mainly sent directly to slaughterhouses, thus reducing their impact on further spatial spread. Beef farms with on-site reproduction and mixed production farms are both potentially more dangerous than fattening farms in propagating the disease because of their transversal and non-trivial patterns of trade movements across different productive classes. As mixed attitude farms are mostly small, the resources required for monitoring them may be limited, and these are the realities that may individually profit most from restoring their full productive potential through disease eradication. However, the use of targeted measures in our simulations predicts that even complete coverage of this productive class would be no better than the random application of these interventions. The relative importance and urgency of measures that are specific to this productive type are probably largely dependent on a limited local context, and their consideration at a national scale may be ineffective.

Our study predicts a total PI prevalence between 1.046% (95% CI 1.042-1.050%) and 2.181% (95% CI 2.176-2.186%) from intermediate to high transmission conditions, respectively, independently of the productive class. These figures are in line with the estimates previously reported in a predominance of studies showing similar prevalence in the range of approximately 0.5% and 2% PI animals in different countries (Houe, 1999), and thus support our choice for the two higher values of the transmission rate explored in this study. Moreover, given that ε leads to an almost linear rescaling of the prevalence values (Fig. 5), our simulation results suggest that a plausible range of values for *ε* may be 0.0012-0.02, based on the comparison with empirical evidence.

Our estimates for within-class dairy farm prevalence (8%) are in line with field observations and modeling predictions on dairy farms in France (6% to 10% (Courcoul and Ezanno, 2010)) and Italy (8.8% (Ferrari et al., 1999)). Direct comparisons with disease prevalence estimates in Italy is difficult, however, since most sources focus on specific areas or single premises or premise types (Cavirani et al., 2013; Nigrelli et al., 2009; Luzzago et al., 1999; Ferrari et al., 1999), thus limiting their generalizability to the national level. Also, the lack of a nationwide control and surveillance plan in the country implies a strong heterogeneity of vaccination patterns and control measures. Given the current situation, our findings may help better informing the design and implementation of a national strategy for intervention and surveillance.

From a modeling point of view, our approach couples within-farm infection dynamics with cattle trade movements at the national scale, accounting for data-driven demography and herd structure. This is an important expansion to the existing modeling literature for this pathogen, as previous works have mainly focused on the within-farm scale, investigating the infection dynamics within the herd structure (Viet et al., 2007; Ezanno et al., 2007; Gunn et al., 2004; Cherry et al., 1998; Innocent et al., 1997b,a; Sørensen et al., 1995; Damman et al., 2015; Smith et al., 2009; Ezanno et al., 2008; Viet et al., 2004), spurred by the intrinsic complexity of BVD disease transmission and the lack of accurate parameter estimates. Importations, however, have been recognized to be important for the maintenance of the disease in the farm (Courcoul and Ezanno, 2010; Ezanno et al., 2008), and synthetic processes for introductions have been considered. Data-driven movements at the national scale were previously integrated in a farm-to-farm transmission dynamics, neglecting within-farm structure and transmission between animals (Tinsley et al., 2012). Neighboring farm effects were considered in a 100-patches metapopulation model for BVD, however trade patterns were assumed to be homogeneous in topology and fluxes (Courcoul and Ezanno, 2010). Several studies have shown on the other hand the large heterogeneity associated to trade movements (Bajardi et al., 2011; Kao et al., 2006; Vernon and Keeling, 2009; Lindström et al., 2010; Dutta et al., 2014). In the Italian dataset for 2014, for example, the number of yearly exchanges with neighboring farms for animal purchases ranges from 1 to 7019 (1-656 for animal sales), and animals are moved in yearly batches of 1 to 1534 units. More importantly, the amount of incoming and outgoing traffic is very heterogeneous across premises, with many of them exchanging few animals, and some hubs featuring several connections to many other holdings (Valdano et al., 2015). Finally, comprehensively considering all productive realities as well as the entire cattle trade system (144,403 premises vs. 100 dairy farms in (Courcoul and Ezanno, 2010)) allows us to capture the entire trade dynamics, where farms of different productive classes are coupled together by non-homogeneous movements, as well as with marketplaces and other types of premises.

The adopted approach presents, however, some limitations. We do not model transiently infected animals to limit the complexity of our modeling approach when considering multiple scales up to the national scale. Their role in the persistence of the disease is still controversial, with modeling studies considering (Viet et al., 2004; Cherry et al., 1998; Innocent et al., 1997a) or not (Gunn et al., 2004; Sørensen et al., 1995) the associated compartment. Our choice may thus lead to lower farm prevalence values than the 33-53% estimated in the literature (Nigrelli et al., 2009; Luzzago et al., 1999; Houe, 1999). These farm-level figures, however, need to be interpreted with caution, given that the definition of infected farm varies widely across studies (e.g. depending on the minimum number of PI, or PIL, or transiently infected animals present in the farm). For this reason, animal-level prevalence reaches a wider consensus.

While we consider the entire geotemporal dataset of trade movements between premises, we do not include pastures, as these are not compulsorily tracked in the national database. Mixed production farms and beef farms are those that may be mostly affected by the lack of these movements, since the practice of moving to pasture in Italy is typical of small, traditional farms. While the role of pastures in livestock disease dynamics is currently the object of preliminary investigations (Palisson et al., 2017) due to lack of data and different national practices, neighboring relationships used in (Courcoul and Ezanno, 2010) to model animal escapes or contacts at pasture were found to only influence epidemic size.

## Conclusions

Through a novel data-driven multiscale modeling approach, we show that farm productive realities play an important role in the dynamics of BVD diffusion due to the their herd composition and the trading patterns they establish. We find that BVD epidemic is dominated by dairy farms, containing the highest number of infected animals and representing the largest contribution to the national farm prevalence. Comparison with available estimates and field observations show that our model captures the heterogeneity of BVD dynamics in a realistic way. Most importantly, our study suggests possible avenues for the implementation of efficient interventions based on the targeted application of control measures to dairy farms. This may contribute to the mitigation and eradication of the disease, and reduction of associated costs. Though the presented work focuses on a single country and a specific disease, the approach can easily be applied to other geographical or epidemiological contexts where trade and herd composition are important element for disease diffusion.

## Funding sources

This work was partially funded by:

- Italian Ministry of Health (www.salute.gov.it) RC-05-2013 “Quantitative methods to identify and calculate risk factors for the BVDV spread across the Italian cattle movements network”
- EC-ANIHWA (Animal Health and Welfare ERA-Net www.anihwa.eu) con-tract no. ANR-13-ANWA-0007-03 (LIVEepi).

